# Benchmarking Molecular Methods of Protein Engineering and Surveying Protein Search Space to Bolster Artificial Intelligence-Based Designs

**DOI:** 10.64898/2026.10.16.759664

**Authors:** Sivakumar Ramamoorthy, Ruby Browning, Joy Brandt, Kirin Tisdale, Alyssa Schultz, Maŕıa Victoria Liendro Dalinger, Nick Hengartner, Daniel R. Tauritz, John F. Beckmann

## Abstract

Proteins are complex macromolecules that drive functionality in cells across all life on Earth. Protein engineering and AI-based designs facilitate exploration of the vast search space. Herein, we explore and compare mutagenic methodologies in combination with fast and efficient phenotypic evaluation of a pink fluorophore (Magenta) to map protein search space and inform AI-based designer algorithms. We binned mutated variants as neutral, deleterious, or catastrophic based on the observed phenotype and fitness measurement (relative fluorescent units, RFUs). Importantly, we found that for a given fluorophore, about 80% of random mutations were deleterious for the artificially selected phenotype resulting in decreased fitness, 20% were neutral mutations, and 0.8% enhanced the artificially selected phenotype. Using error prone PCR (epPCR), we were able to identify four candidates in the first round and one candidate in the second round of epPCR for enhanced fluorescence and in toto enhanced RFUs by 35% after two rounds of epPCR. We compared these rates of true positive enhanced RFU generated sequences against a state of the art evolutionary algorithm approach and found that biological computation beats it in its current state. We conclude with suggestions for key improvements to enhance algorithm performance in the future. Taken together, we demonstrated that iterative rounds of variant prediction improved our algorithm, and our study can serve as a reference point for future AI driven modeling studies in red fluorescent protein engineering.

## 1 Introduction

Proteins are organic macromolecules that have evolved over billions of years which power diverse cellular functions across the tree of life. The primary amino acid sequence coding the protein directly influences its function [1]. Enhancing protein functionality for desired applications is a main challenge of modern biotechnology and AI. Protein engineering involves tinkering with the natural sequence to optimize for particular characteristics supporting existing or new functions, such as optimizing proteins for MRI analysis [2]. Creating variants with enhanced fitness quickly is not an easy task [3, 4]. Natural selection gradually evolved the proteins that exist today over billions of years [5, 6]. To engineer proteins more quickly, two discrete pre-AI approaches, directed evolution (DE) [7, 8, 9, 10, 11, 12, 13] and computational protein design (CPD) [14, 15, 16, 17, 18, 19, 20, 21, 22] were employed in parallel and recognized by the Nobel prize committee in 2018 and 2024 respectively [3].

In DE, the process of natural evolution is expedited through iterative rounds of mutation followed by wet-lab artificial selection where each iteration is a step in evolution. Variation inducing strategies are either random, like error-prone PCR, or targeted, like site-directed mutagenesis [23, 24]. Random mutagenesis strategies generate point mutations leading to highly diverse sequence libraries derived from a source gene. These approaches include cell-based, chemical, UV-based, and PCR-based methods [23]. Cell-based mutagenesis involves passing the gene of interest through a host mutator strain possessing faulty DNA repair or DNA replication machinery. Other methods use chemical mutagens, like the alkylating agent N-methyl-N’-nitro-N-nitrosoguanidine (NTG) or UV radiation, which modifies DNA either *in vivo* or *in vitro*. Major downsides of these approaches are that mutations can strike throughout the whole plasmid and host chromosome; they are not specifically restricted to the gene of interest. These strategies also generate epistatic interactions (where effects from one gene modify another’s) and pleiotropic effects (where one gene modifies multiple phenotypes) [25], complicating phenotypic analysis.

Error-prone PCR (epPCR) is one widely used *in vitro* DE method to turbo-charge random mutatage-nesis of a specific target gene [26]. epPCR avoids epistatic effects and generates diverse mutant libraries at a target gene only. This method uses a low-fidelity Taq DNA polymerase combined with buffer and conditions that promote introduction of random mutations during PCR amplification [27]. After mutagenic amplification, the PCR amplicon is ligated to plasmids, creating a genetic library of sequence variants. The mutant library is then transformed into an organism to search the protein sequence space by evaluating phenotypes. Useful variants can be isolated and assessed for genotype-phenotype relationships. The bottleneck of this approach is that it generates large numbers of mutations in the target gene making phenotypic screening of the library laborious, time consuming, and even impossible for some proteins. The libraries generated by epPCR contain a majority of A to G and T to C transitions resulting in technically non-random (biased) distributions of amino acid changes in the given sequence that are generally deleterious [28]. Interestingly, these mutations navigate through a codon hamming graph, which acts as a biological search algorithm traversing through protein space. A key point is that these methodologies are not random.

CPD approaches, in contrast, rely entirely on predicted biophysical models trained by statistical methods and informed by data from the protein data bank (PDB) [29]. Early CPD frameworks like Rosetta operated based on this principle of using a physics-guided energy function and intuitive sampling models. These classical designs operated by searching conformational space to determine sequences that reduce free energy for a given target structure, in theory making it more stable [30, 31, 32].

The fundamental limitation with DE and CPD approaches is that both strategies search through the unimaginably vast space of protein primary structures. A small protein comprising of only 100 amino acids generates 20^100^ *∼* 10^130^ protein sequence variants, far surpassing the estimated total number of matter particles in the universe (4*x*10^80^) [33]. Moreover, only a minute fraction of these sequence variants can fold and are functional. In principle, both DE and CPD approaches function by navigating through the protein search space with functional validation of variants. To search the space, DE relies on the experimental scoring of variants directly measuring the desired properties, while the promise of CPD is to scan further into the protein search space; but this approach suffers from the drawback that it relies on a single energy function irrespective of the protein’s unique phenotypic functions. To overcome this, these two strategies are often used in conjunction to complement each other such that data obtained from DE will be used to inform and refine CPD variants [34, 26]. The limitation of machine learning (ML) guided directed evolution models is that the functional predictions depend on limited training sets that are biased; and structural folding algorithms must correctly model amino acid dependencies throughout the whole protein. A single amino acid change in one position can impact and alter the function of that protein and even other proteins [35]. The failure to model or train these epistatic effects results in inaccurate predictions of fitness [36, 37]. Collectively, these limitations lead to a nongeneralizable approach, where execution is inefficient and often protein specific.

In an attempt to overcome limitations of ML guided directed evolution, we leverage the Evolutionary Algorithms Simulating Molecular Evolution (EASME) package we have developed, that models DNA strings within an evolutionary algorithm (EA) integrating triage filtered fitness functions driven by bioinformatics. EAs are heuristic algorithms mimicking natural evolutionary processes. These algorithms simulate evolution by coupling stochastic processes of substitution, mutation and recombination to explore sequence space. Each mutated sequence is then assigned a fitness value that are used to emulate selection. Our hypothesis is that EAs improve upon other forms of AI/ML, as EAs are particularly good at traversing complex fitness spaces and do not necessarily require training data as a reference. Starting with a protein sequence, the algorithm effects mutations and recombinations to produce other sequence variants [38]. EASME is powered by bio-inspired AI that represents a paradigm shift in how biological problems are approached using AI on computers. This opens new research opportunities in the fields of biological and computer science by introducing an independent, complementary, synergistic approach to ML and large language model (LLM) designs. Rather than converging upon predetermined structural targets limited by training sets, EASME uses biologically informed constraints to navigate and explore DNA/protein sequence space, collectively unlocking novel capabilities just as biology does. Our group previously modeled the evolution of *Wolbachia* specific toxin-antidote systems responsible for reproductive parasitism in arthropods by employing EAs to simulate evolution of actual DNA strings [39]. This study laid the foundation to extend the approach for bio-engineering any given protein sequence.

To better inform the design and assessment of AI-based protein engineering algorithms, we benchmarked wet lab mutagenesis procedures to compare efficiencies and roughly map protein search space of a pink fluorophore, Magenta, with the intent of artificially selecting better fluorophores and designing better AI de novo protein generators through EASME. For clarity, we follow biological naming conventions defining Magenta as the protein, *magenta* as the gene, and pMagenta as the plasmid encoding the gene. We chose fluorescence as our artificially selected phenotype because (i) it is easy to measure, (ii) enhancing the fitness is difficult, and (iii) fluorophores are widely engineered proteins [40]. Thus, fluorescent proteins have been engineered for decades and represent a useful starter model to benchmark comparative AI models against and ground them in biological truth. In other studies, ML models and high-throughput wet lab screening yielded improved green fluorescent protein (GFP) sequences with enhanced fluorescence, incorporating 40 to 50 mutations in 238 amino acid coding sequences [40].

In this study, we directly compare efficiencies of truly random site directed mutagenesis, UV mutagenesis, natural variation, epPCR, and finally an initial EASME prototype design, in exploring protein search space of fluorophores. Specifically, our selected phenotype was to simply enhance relative fluorescent unit (RFUs) of Magenta. Through our experiments, we were able to directly estimate rates of neutral, deleterious, and beneficial mutations given an artificially selectable laboratory phenotype, providing useful benchmarks. We map mutations detrimental to, or improving RFUs, and test current algorithmic methods using EAs. Overall, we successfully incorporated the data obtained from conventional approaches to train an EASME mutational operator in a design-build-test-learn (DBTL) feedback loop, that moderately improved algorithmic navigation through the protein search space. Importantly, we compare in laboratory driven evolution against *in silico* evolution. Our results demonstrate that the EASME pipeline can be trained and can improve, invigorating future exploration of this novel bio-inspired algorithmic approach.

## 2 Materials and Methods

### 2.1 Bacterial strains and growth condition

The bacterial strain and plasmid constructs used in the study are listed in Table S1. *Escherichia coli* Top 10F’ was routinely sub-cultured in lysogeny broth (LB) medium, grown at 37°C with agitation at 200 rpm. When needed, the medium was solidified with 2% agar. Antibiotics were added as required at the following concentration (unless otherwise specified): ampicillin (Amp), 100 µg ml^-1^; chloramphenicol (Cm), 34 µg ml^-1^.

### 2.2 Optimization of bacterial cell density and time for RFU measurement

We performed an initial time course experiment to normalize the optimal bacterial cell density and time of growth for fluorescence measurement. Since we observed a difference in phenotype after reviving the stock, we inoculated individual colonies displaying bright, intermediate, and mild pink fluorescence into LB medium supplemented with Cm. The culture was grown for 48 and 72 hours. The cell density was adjusted to 0.5, 1.0, 1.5 and 2.0 at *OD*_600*nm*_ and the fluorescence was measured using QFX fluorometer (DeNovix, USA). Normally, the Y axis represents the relative fluorescent units (RFUs) and X axis the optical density (OD) at 600 nm. The experiment was performed in triplicate and the result was represented as mean with standard deviation (Figure 1A).

**Figure 1:**
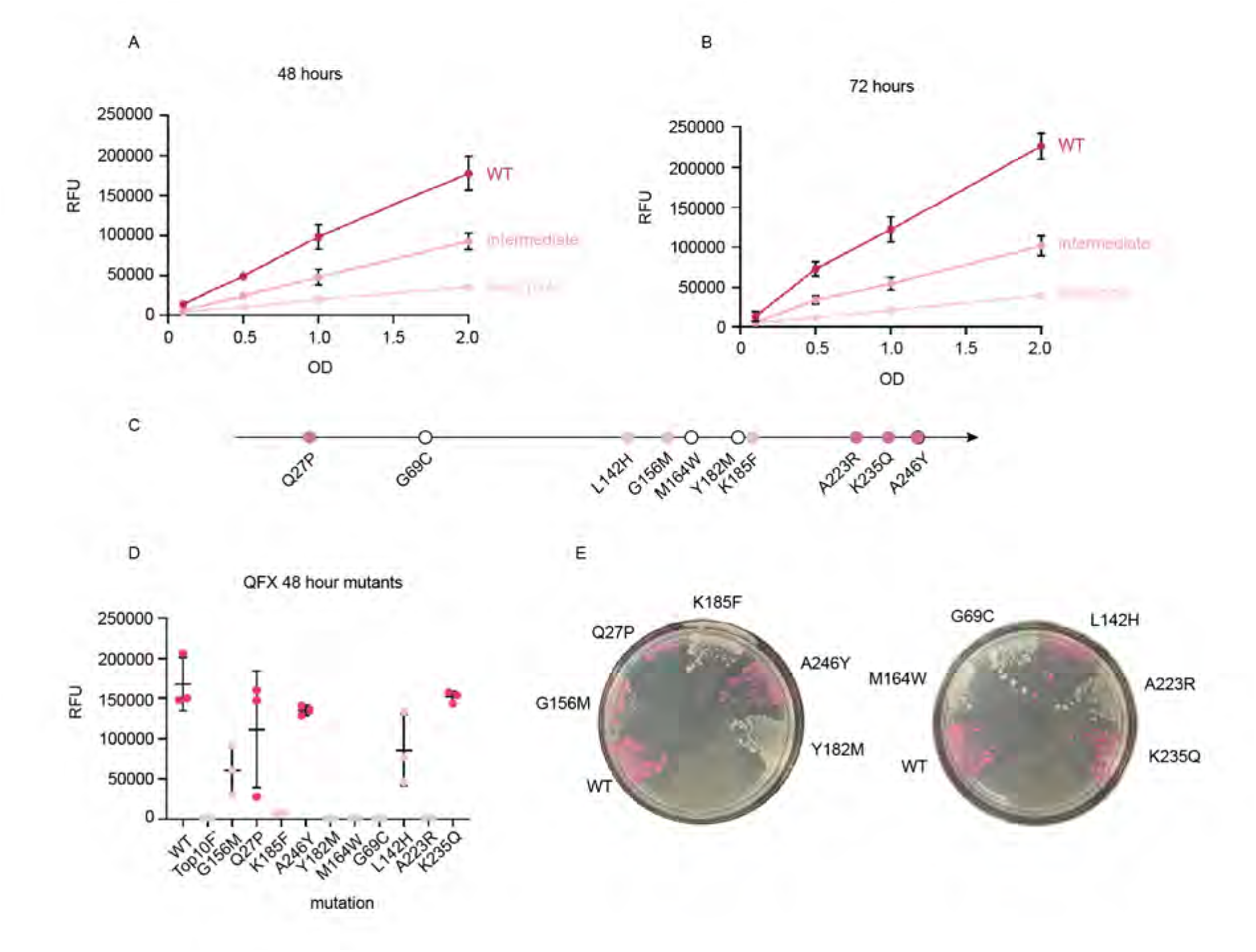
Calibration of fluorometer experiments and random walk mutagenesis. **A.** 48-hour analysis of WT pMagenta variants. Three clones are analyzed with most pink WT, intermediate pink, and least pink shown. **B.** Similar graph as in A., at 72 hours time point. **C.** Mutation map of mutations derived from a random walk mutagenic algorithm. Pink circles are neutral mutants with most pink fluorescence, intermediate pink mutants are deleterious mutations, and white circles are catastrophic breaking mutations eliminating fluorescence. **D.** Quantified QFX fluorometer data of the random walk mutations. **E.** Streak plates of random walk mutations correlates mutant phenotypes with RFUs in D.

### 2.3 Measuring RFU with QFX fluorometer

Starter cultures of Top10F’ *E.coli* harboring pMagenta were inoculated in 3 mL LB with Cm and allowed to grow to saturation for 48 hours. Cultures were standardized to a final concentration of 1 OD in 200 µL via spectrophotometry at 600 nm. RFUs were measured in a QFX fluorometer (DeNovix, USA) using the green excitation channel, measuring output at 565-650 nm excitation (Figure 1A).

### 2.4 Random Walk Mutation Algorithm and Site-Directed Mutagenesis

Magenta has 263 amino acids (see Figure S1). A random dice roll from 1-263 was conducted 10 times to pick random amino acids to mutate. Amino acids selected by this algorithm were 27, 69, 142, 156, 164, 182, 185, 223, 235, and 246. Subsequently, each of the 20 standard amino acids were assigned a number from 1-20, and a 20-sided dice was rolled to pick the amino acid to mutate to. After mapping of the mutations, site-directed mutagenesis primers were designed and polymerase chain reaction (PCR)-based mutagenesis was performed on the pMagenta according to standard methods [41, 42]. Mutated DNA was transformed into *E. coli* Top10F’ by classical chemical transformation, colonies were selected and the mutants were confirmed by sequencing.

### 2.5 UV Mutagenesis

*E. coli* Top10F’ strain harboring the pMagenta was struck on LB medium containing Cm antibiotic from frozen stock. A colony was inoculated to LB broth and grown at 37 °C with agitation for 48 hours. Serial dilutions of the culture were made and 10 µl of 10^4^, 10^5^ and 10^6^ were plated on LB medium containing Cm antibiotic. The plates were immediately placed inside a bio safety cabinet and exposed to UV light for 7.5 s. Longer exposure to UV light resulted in complete killing of the bacteria. After exposure to UV light the plates were incubated at 37 °C for 48 hours. Clones were re-struck and categorized as bright, intermediate and white clones (Figure 2A,B,C). Ten representative clones from each category and unmutated *E. coli* harboring pMagenta plasmid, which served as control were inoculated into 5 ml LB broth with antibiotic and grown for 48 hours. Post 48 hours of growth the *OD*_600_ nm was adjusted to 1.0 and the fluorescence was measured using QFX fluorometer as described above. To identify the mutations plasmids were extracted using E.Z.N.A mini extraction kit (Omega Bio-tek, USA) and whole plasmid sequence was ascertained at Plasmidsaurus sequencing facility. Previously, genetic screens of UV induced mutations in the *lacI* gene of *E. coli* revealed 60-65% of mutants showed base substitutions, 30 to 35% showed frameshifts while 5% of the mutants displayed deletions of several bases [43]. Our hypothesis was UV mutagenesis would induce mutations in *magenta* improving fitness.

**Figure 2:**
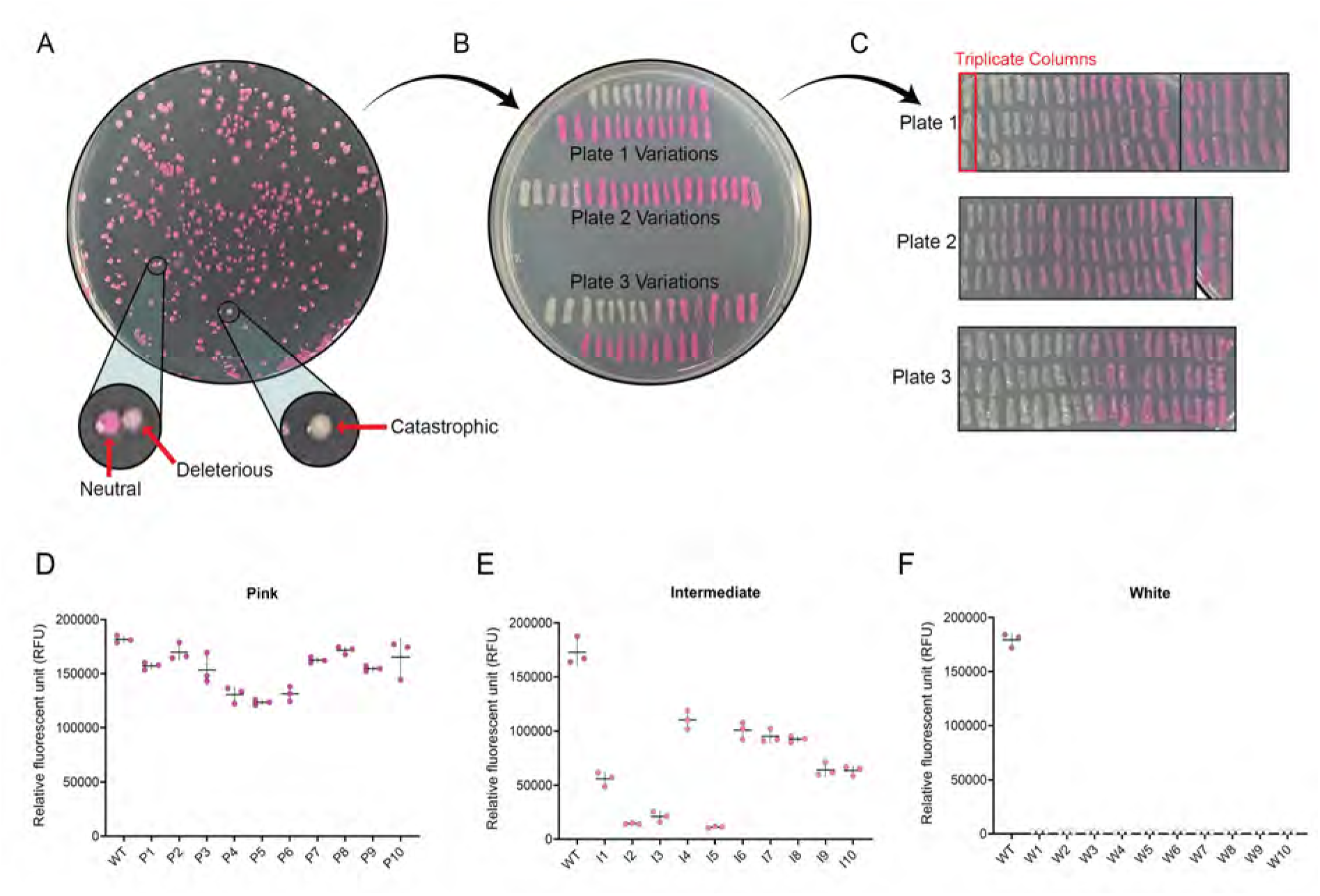
UV mutagenesis of *E. coli* expressing Magenta for purposes of artificial selection. **A.** Plate of *E. coli* colonies exposed to UV light for seven seconds. Pink colonies express Magenta. Circles and red arrows point to mutants that are either slightly deleterious (intermediate pink, left) or catastrophic (white, right). Plate is representative of biological triplicates. **B.** Streak plate of picked colonies from A. representative of phenotypic variants observed including white knockouts, intermediate pink deleterious mutants, and wildtype or neutral clones. **C.** Triplicate re-streak of colonies from B. to confirm phenotype consistency. Verticals indicate triplicates of the same clone. **D.** RFU of ten most pink clones derived from the UV-mutagenesis experiments. WT is wildtype control and P represents pink colonies. **E.** RFUs of ten intermediate (I) clones. All clones show significant reductions in fluorescence when compared against the WT control. **F.** RFUs of ten catastrophic white (W) mutants.

### 2.6 Error-Prone PCR

Earlier, several groups used epPCR successfully and synthesized libraries with 15 to 30 nucleotide mutations per gene [44, 45, 46, 47]. To map the beneficial mutations responsible for the enhanced fluorescence we set out to generate random mutant library using epPCR as described previously with modifications [48]. Briefly, epPCR was performed using pMagenta as template. The PCR reaction was carried out using primers targeting the pink gene and Taq polymerase (New England Biolabs, USA) owing to its natural high error rate [49]. The list of primers used in the study is shown in Table S2. To increase the error rate, we used *MgCl*_2_ and *MnCl*_2_ at concentrations 5 mM and 0.25 mM respectively. The PCR conditions included one cycle at 94 °C for 10 min, followed by 40 cycles at 94 °C for 30 s, 55 °C for 30 s, and 72 °C for 90 s, with a final elongation step at 72 °C for 5 min. The generated library was resolved on 1% agarose gel, and the amplicon was eluted using E.Z.N.A gel extraction kit (Omega Bio-tek, USA) following manufacturer’s instructions. The eluted product was ligated to pGEM-T easy (Promega, USA) using T4 DNA ligase (New England Biolabs, USA) following manufacturer’s protocol. The ligated product was transformed into *E. coli* Top10F’ chemically competent cells using the heat shock method [50]. The transformants were plated on LB medium containing ampicillin, IPTG 40 µg ml^-1^, X-gal 20 µg ml^-1^ and plates were incubated at 37°C for 16 hours. The unmutated gene coding for the pink protein was cloned into pGEM-T vector and served as wild type (WT) control, since a significant difference in fluorescence was observed between pMagenta parent plasmid and WT pGEM-T clone (Figure 3A). To further explore the search space, we performed iterative rounds of epPCR using an enhanced clone from round one as template (mirroring a biological hill climb with each cycle one step further in the evolutionary chain). To dissect the mutations responsible for enhanced fluorescence of positive clones obtained via epPCR, we performed site-directed mutagenesis as mentioned above. All experiments were done in triplicate and results were expressed as the mean with standard deviation.

**Figure 3:**
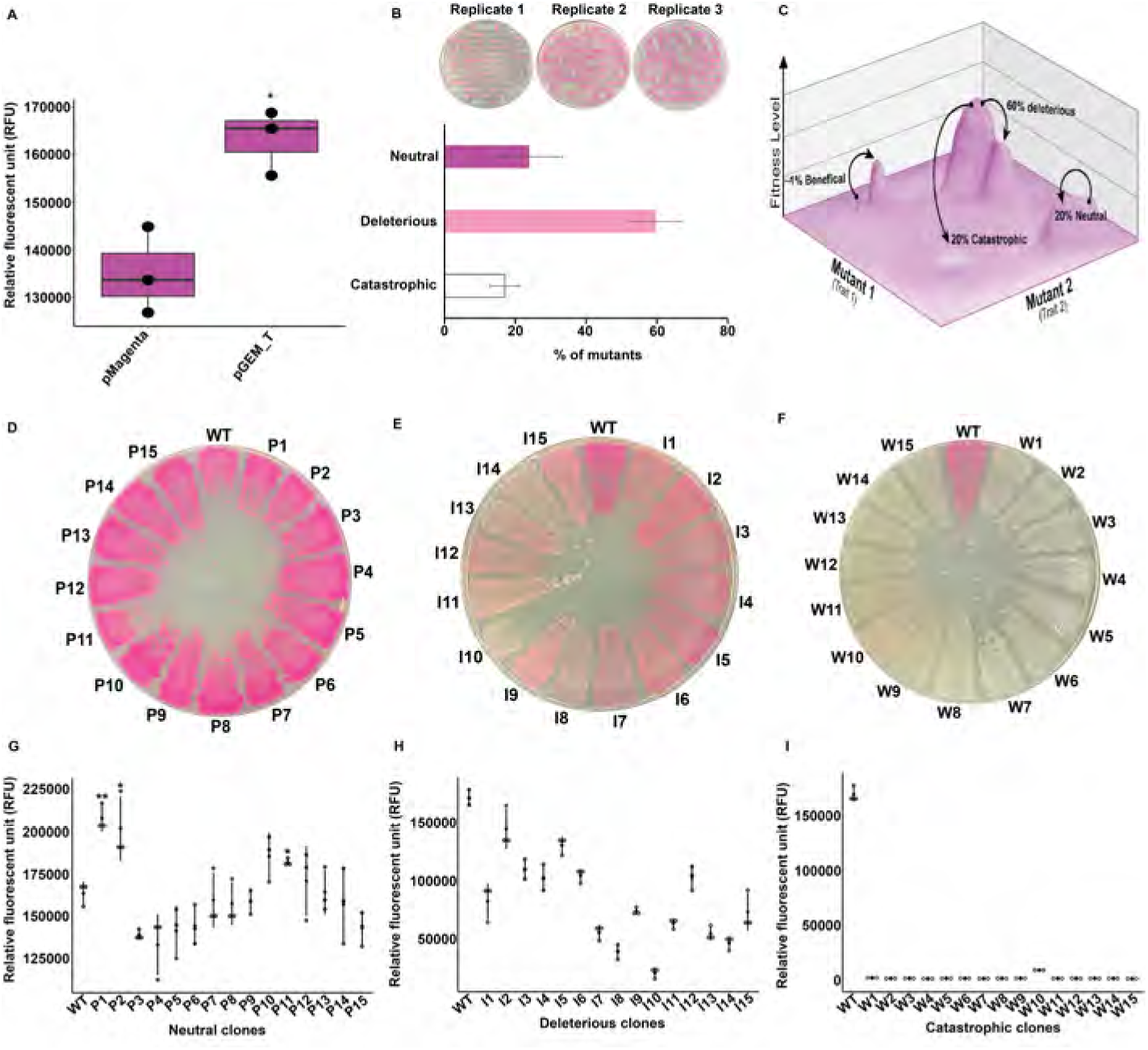
Scanning the search space of pink fluorophore for beneficial mutations using epPCR. **A.** Box plot showing the difference in fluorescence between the parental pMagenta and pGEM-T plasmid, used to construct mutant library (*, p*<*0.05). **B.** Graph showing the percentage of neutral, deleterious and catastrophic mutants generated (Below) and representative plate image of the mutants (Top). The experiment was performed in triplicate. Results were represented as mean with standard deviation. **C.** Abstract fitness landscape showing mutant proportions. **D.** Representative plate image showing neutral mutants relative to WT. **E.** Representative plate image showing deleterious mutants in comparison to Wild type. **F.** Representative plate image showing catastrophic mutants in comparison to WT. **G.** RFU measurement of 15 neutral mutants (*, p*<*0.05) (P represents pink colonies). **H.** RFU measurement of 15 deleterious mutants (I represent intermediate pink colonies). **I.** RFU measurement of 15 catastrophic mutants (W represent white colonies). RFU is measured in triplicate as mean with standard deviation.

### 2.7 EASME Approach

EASME is a new sub-field of evolutionary computation [51], the field that studies EAs. While evolutionary computation is often employed to solve engineering problems [52], EASME aims to use evolutionary computation to solve *biological evolution* problems, allowing for entirely *in silico* proteomics pipelines. One of the goals of this project was to benchmark lab-based protein engineering methods against a digital EASME prototype, in order to identify problems and debug them. The EASME package can be used to implement biologically informed AI for bio-engineering proteins. Two of its core algorithms are (1) the exploration of protein sequence space, and (2) model-based evaluation of fitness. Both algorithms can be adapted to the specific bio-engineering problem being solved, possibly in an iterative manner that incorporates experimental results as they become available.

#### 2.7.1 Data Used

The first round of EASME mutants was designed using an EA evaluating fitness through comparison with 118 proteins downloaded from the UniProtKB database. A BLAST search was performed on the original Magenta protein, with the most similar proteins downloaded for our data set. These proteins were split into two groups based on markers in their descriptions — “fluorescent” (58 proteins) or “nonfluorescent” (60 proteins). The EASME package used the amino acid sequence of pMagenta as a starting point, then produced numerous mutants, attempting to maximize homology with the fluorescent data while minimizing homology with the non-fluorescent data. Fitness of mutated variants was evaluated by homology against the two sets of proteins. Higher alignment scores with close fluorescent homologs was rewarded, while higher alignment with the non-fluorescent proteins was penalized. The homology score was evaluated by averaging pairwise homology between the target protein and every other protein in each of the two reference groups.

For the second round, mutation positions were *not* selected entirely at random, but using a *weighted* random sample. Twenty random fluorescent proteins were chosen from the entirely-fluorescent FPbase [53] data set (to reduce the computational time required for sequence alignment evaluations), then a multiple sequence alignment was performed comparing them to the pMagenta sequence. The rate at which each position was found to be divergent from amino acid sequence of the pink fluorophore protein determined the likelihood that that position would be chosen for mutation — i.e., more conserved regions of the protein were less likely to be mutated. ESM’s public ESMFold API was used to produce a folded 3D model of the new variants, and the distance between the chromophore and the center of the protein was evaluated as a final measure of fitness (ideally, the chromophore should be located in the center of the protein’s “beta barrel” structure [54]).

Following only minor improvement in the second round, a more comprehensive data set was desired. We downloaded the FPbase, an online database specifically for fluorescent proteins, featuring relevant data such as excitation wavelengths and emission wavelengths (which together contribute to color and RFU of a fluorescent protein). For the purposes of this project, we set a target excitation value of 542.5 nm, and a target emission value of 620 nm. Fluorescent proteins closer to these target values were given proportionally larger preference in the fitness function — homology to another red protein was scored higher than homology to a green protein, for example. This fitness function was assessed by whether it correctly evaluated the random walk mutants generated in the wet lab.

#### 2.7.2 *In Silico* Mutagenic Process

The first round utilized fairly relaxed mutation parameters — each generation, any protein in the population had a 12.5% chance of being duplicated and then mutated, and mutations were entirely random — a random choice was made between insertion, substitution, and deletion, a random length between one and the length of the protein was generated, a random start point was selected, and in the cases of insertion and substitution, random new subsequences were generated from the 20 standard amino acids. This approach produced several variants that were quite different from the original protein, but as can be seen from the first round data in Figure 6, none of these ended up being functional.

The second round utilized a much more conservative mutation function, mutating only set numbers of amino acids, then never modifying an already-mutated variant again. Mutation rates were one amino acid, three amino acids, and five amino acids respectively — unique positions would be selected, and then each amino acid at those positions would be replaced with a randomly-chosen amino acid.

The third round, like round two, aimed to mutate the start sequence such that it aligned better with other proteins closer to the target excitation and emission values. This round produced better results, though these may be attributed to the fact that it was discovered between rounds that the previous two utilized an accidentally truncated start sequence. It is interesting to note that, even with part of the sequence missing, the second round variation operator possibly restored some fluorescent functionality.

### 2.8 EASME Clones Construct Assembly

An EASME algorithm was written (see Section 3.5) and used to generate three rounds of 15 starter sequences to test, for a total of 45 new sequences. Sequences derived from EASME were DNA codon optimized for *E. coli* tRNA preference and ordered via GenScript synthesis into pUC57 backbones with flanking 5’ BseRI and 3’ SacII restriction endonuclease sites. The 15 clones were excised via restriction digest with calf intestinal phosphatase and subcloned into the pMagenta backbone with endogenous promoter replacing the original WT. Clones were transformed as above and screened via Sanger sequencing. Additionally, promoters were sequenced to validate that the promoter was perfect. For the second and third rounds the generated variants were order via GenScript with the native promoter and synthesized into pUC57. The plasmid was transformed into *E. coli* Top10 F’ and fluorescence was measured on the transformants.

### 2.9 Statistical analysis

Data were generated from one independent experiment performed at least in triplicate, and the results are represented as mean*±SD*. All statistical analysis were performed using R. Statistical significance between wild type and the variants was evaluated using one way ANOVA with Tukey’s post hoc test for multiple comparisons (for normally distributed data) or Kruskal Wallis test (a non parametric test) when the Shapiro–Wilk test failed to support the normality assumption. For all analysis the null hypothesis of no difference was rejected at the p*<*0.05 level.

## 3 Results

### 3.1 Exploring Molecular Methods of Mutagenesis for Artificial Selection

Our hypothesis was that we can utilize modern methods of mutagenesis to create variants that can be artificially selected in the laboratory to produce both higher intensity pinks and enhanced RFUs. In effect we were creating a biological EA. We also wanted to discern the best molecular variation operator for mutagenesis and sought to compare methods to tune the biological computation. To begin artificial selection on Magenta, we first optimized a fast approach to assess its phenotype from expression with pMagenta transformed into *E.coli* Top10F’ bacteria (see Figure 1A and B). Because the fluorophore expresses within bacteria, the RFUs measured directly correlate with bacterial cell concentrations and growth curves. To standardize the process for all future experiments we compared RFU measurements performed at 48 and 72 hours of culture, measuring concentrations of bacterial cells at OD 600 nm.

For WT constructs, RFUs accumulate and saturate around 200,000 RFU at 72 hours (see Figure 1B). Thus, we concluded that the optimal time to discern differences in RFU counts per cell were at 48 hours growth. This protocol left ample measurement room to detect increases and decreases in RFUs after mutagenesis and artificial selection. From this growth timeline we were able to visually observe and discern differences in clonal population RFUs for *≈* 3 distinct bin categories of pink shades (pink (WT), intermediate pink, and least pink). After determining the process for optimal RFU measurement we began to explore mutagenesis methodologies with the goal of artificially selecting higher RFU’s/cell.

Of the various processes to generate pseudo-random molecular mutagenesis listed in Section 2.7.2, we explored three simple initial methods as *variation operators* to generate mutations in the gene coding for *magenta*, with intent to find the best methodology for searching protein space for a fluorophore delivering higher RFU/cell. Biology itself does not have a completely random mutation process, so we designed a simple non-biological random mutator described in methods Section 2.7.2 and executed 10 truly random mutations via site directed mutagenesis (see Figure 1C, D, and E). Of the ten mutants constructed, 40% were catastrophic mutations that destroyed the fluorophore, 40% were moderately deleterious, and 20% were neutral. None of the ten were beneficial in terms of our artificially selected phenotype, enhanced RFUs/cell. Importantly, this process was cumbersome.

### 3.2 UV mutagenesis was not an appropriate strategy to scan the protein search space

Having established an initial baseline rates for neutral, deleterious, and catastrophic mutations, we tested other molecular *variation operators* to discern if we could improve upon these rates; we tested hunting for natural variation mutants in large populations and UV mutagenesis. Mutations in *E. coli* occur approximately at rate 2.2 *×* 10*^−^*^10^ mutations per base pair per generation [55]. This makes plating large numbers of clones impractical to generate and observe mutant diversity in pMagenta. This was confirmed experimentally, with clonal colonies showing essentially zero mutants (Figure S2).

Next we explored UV mutagenesis as a variation operator. As expected, UV mutagenesis generated more visible mutants than natural variation. We could not quantify rates of deleterious/catastrophic mutants because it was impossible to ascertain the denominator. Another negative aspect of UV mutagenesis is that it generates non-specific mutations not necessarily in the gene of interest. While it produced more variants than natural evolution, it was unable to produce an enhanced RFU mutant (see Figure 2). To assess fluorescence of mutants, we generated UV mutants on plates, struck clones out in triplicate, and quantified RFUs in triplicate for 10 neutral, 10 deleterious, and 10 catastrophic mutant clones. We also sequenced their plasmids (discussed below). None of the clones we assessed improved the RFUs/cell (n=30) (Figure 2D, E and F).

The sequencing of the ten mutants with catastrophic deleterious mutations that resulted in the complete loss of fluorescence, revealed that three mutants (Mutant W2, W6 and W9) had deletions in the promoter, three mutants (Mutant W1, W3 and W5) had insertion in the coding region and four clones (Mutant W4, W7, W8 and W10) had no insertions and deletions (i.e., indels) in pMagenta (Figure S3A). We reasoned that the loss of phenotype in these mutants can be a result of chromosomal mutations or other nonspecific mutations on the plasmid backbone. To validate our hypothesis, we re-transformed plasmids extracted from mutants W4, W7, W8 and W10 to *E. coli* Top10’ and all transformants appeared white (Figure S3B). Analysis of sequence revealed insertion of transposon element was observed in the plasmid backbone of W4, W8 and W10 mutants while in Mutant W7 recombination of plasmid backbone was observed resulting in a large indel (Figure S3C). Of the sequences obtained, all ten deleterious mutants (less fluorescing mutants) showed no changes in the gene and plasmid sequence, suggesting that UV mutations had struck the *E.coli* chromosome causing epistatic and pleiotropic effects that had an impact on RFUs/cell without changing the actual fluorophore gene itself. To test this hypothesis, we re-isolated the plasmids and re-transformed them into fresh Top10F’ *E. coli* cells. After transformation, the color reverted back to wildtype, supporting the conclusion that UV mutagenesis was generating pleiotropic/epistatic effects (Figure S3D and E). Taken together, our data indicates that UV mutagenesis is not a useful approach for artificial selection of proteins as it induced large scale deletions and transpositions indels in our hands. Thus, pure random mutation, natural variation, and UV mutagenesis failed initial feasibility tests for variation operators in a biological EA. Thereafter, we switched focus to epPCR.

### 3.3 epPCR is the Best Mutagenic Operator for Artificial Selection

To address the lack of improvement in fluorescence phenotypes using prior methodologies, we chose to experiment with epPCR. In contrast to the other methods, epPCR specifically targets a gene of interest for mutation. We adjusted the manganese (*Mn*^2+^) and magnesium (*Mg*^2+^) ion concentrations of PCR reactions targeting *magenta* to 0.25 mM and 5 mM respectively; these concentrations specifically induce errors in PCR. Our hypothesis was that this methodology was a better molecular search algorithm in protein space.

Thus, we generated mutant libraries of pMagenta and quantified their associated RFUs as our artificially selected phenotype. This enabled us to quantify and compare whether epPCR explored the sequence space more efficiently than other methods that do not target the gene of interest specifically. Simultaneously we aimed to both enhance RFUs and map the fitness landscape of *magenta*. In total, we generated triplicate independent epPCR mutagenesis libraries targeting the gene and promoter sequence for magenta. The resulting clones from each library were binned by eye into three categories, including pink (neutral), intermediate (deleterious) and white (catastrophic) mutants (n = 1250). From these samples we calculated the ratio of different mutation bins revealing 80% of epPCR mutants generated were either deleterious or catastrophic (Figure 3B & C). 20% of mutations were neutral and a small subset of those were beneficial. Importantly, epPCR was the only methodology capable of producing beneficial mutations that we could recover en masse.

To identify beneficial mutations, we quantified RFUs from a sample of 100 randomly selected neutral clones from first round of epPCR. Four clones (P1, P2, P10, and P11) showed enhanced RFUs when compared to the wild-type plasmid (Figure 3D and G). 20% of the total library were binned as neutrals— clones that are observably as pink as WT (Figure 3B). Of those, 4% were actually beneficial. Thus, the rate of beneficial mutants (and the rate of true-positives from these biological computations) produced from epPCR was roughly 0.8% of the total mutants produced. In addition to neutral clones, we also measured the relative fluorescence of 15 deleterious (Figure 3E and H) and 15 catastrophic mutants (Figure 3F and I). At the moment, this is our current state-of-the-art rate of finding better variants for an artificially selected phenotype using a biological EA. A researcher can experimentally inflate this rate to *≈* 4%, if the protocol can quickly sort neutrals from deleterious, as we show. Importantly, this number reflects a state-of-the-art true-positive rate that AI based protein engineering methodologies should set goals to beat in order to outperform modern molecular methodologies themselves. Biological researchers themselves should seek to boost these values.

To map the spatial distribution of mutations on the plasmids of clones showing enhanced RFUs, we sequenced and BLAST compared the four enhanced RFUs clones P1, P2, P10 and P11, against WT. Mutant P1 had two amino acid substitutions, M2I and N249S and a synonymous mutation resulting in no amino acid change. Mutant P2 had one coding mutation, T255A, and a transition mutation in the promoter sequence -248A*>*T. In Mutant P10 we identified a promoter transition mutation (-3T*>*A) resulting in insertion of an upstream start codon (methionine) and a mutation in the coding sequence S248G. Mutant P11 had a single transversion mutation in the promoter region -36A*>*G. This methodology is not random, *Taq* polymerase has a natural tendency to introduce certain types of errors over others [56]. For example, they are highly biased towards transitions over transversions, which was evident in our study. We observed 75% transition mutations in contrast to 25% transversions (Figure 4A). Based on the sequencing results, on average two mutations were introduced in the gene. Because enhanced RFU resulted from mutations in either the coding or promoter regions, we suggest that bio-engineering protocols will need to distinguish between improvements in the gene function and changes in gene expression levels.

**Figure 4:**
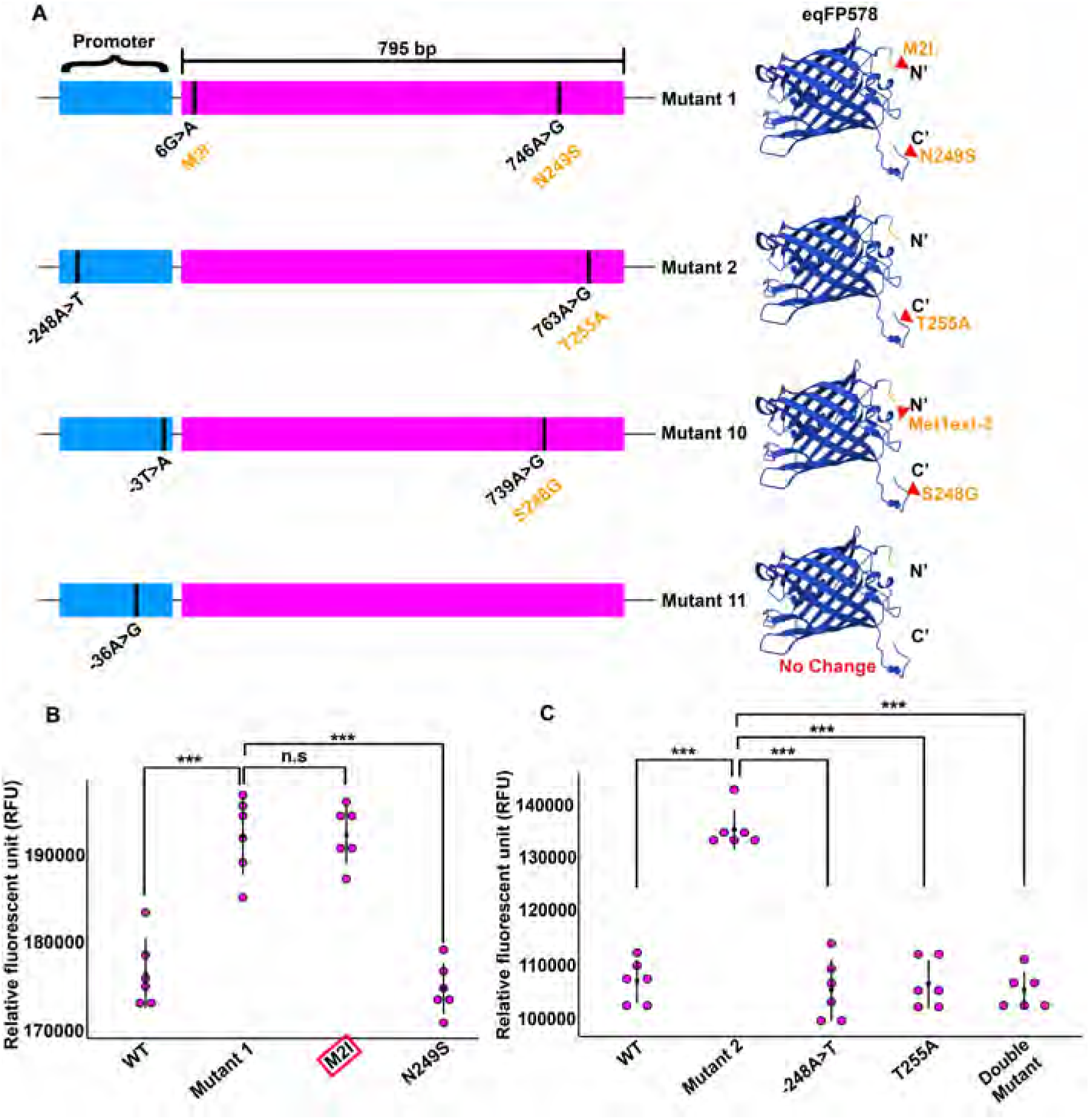
Mapping of mutations in clones exhibiting enhanced fluorescence relative to the WT. **A.** Graphical representation of genetic organization of gene coding for Magenta. The black line represents the position of mutation on the gene, and the corresponding amino acid substitutions are denoted below. On the right is the protein structure predicted by AlphaFold 2, showing the position of amino acid substitution. Site directed mutagenesis to dissect the amino acid substitution was involved in increased brightness of Mutant 1 **(B)** and Mutant 2 **(C)** (***, p*<*0.0005; n.s, not significant).

When mutations compound, it is hard to discern which individual mutation, or synergistic combinations, cause the observed change in phenotype. To attribute the increase in fluorescence of Mutant P1 and Mutant P2 to individual mutations, we performed site directed mutagenesis (SDM) to isolate and test each mutation to quantify its singular impact on RFUs. In Mutant P1, SDM results revealed a single amino acid substitution M2I was responsible for the increase in fluorescent phenotype (Figure 4B). In Mutant P2, neither promoter mutation nor amino acid substitution at position T255A alone was responsible for enhanced fluorescence, leading us to hypothesis of a synergistic effect of both mutations. To test this, we constructed *de novo* a double mutant from the neutral clone harboring only these two mutations. This controlled double mutant displayed significantly lower fluorescence when compared to Mutant P2. This suggests that P2 contains one or more epistatic mutations, indicating a possible different mechanism for increasing the RFU in Mutant P2 (Figure 4C). For context, the tertiary structure of most red fluorescent proteins are highly conserved with the chromophore region sharing a tripeptide sequence (X-Tyr-Gly) inside a capped beta barrel [57, 58]. Red chromophore maturation results from a chemical conversion of GFP-like chromophore that is protonated [58]. We speculate plasmid backbone mutations could be responsible for the enhancement in RFU. Further investigation of Mutant P2 is needed to determine this.

### 3.4 Iterative epPCR Further Enhances Artificially Selected Phenotypes

Having validated that epPCR can produce variants for artificial selection of RFUs, we investigated if additional application of epPCR followed by phenotypic selection leads to further improvements. We subjected Mutant P1, which contained two amino acid substitutions in the coding region, to a second round of epPCR random mutagenesis. We screened the generated library for increases from B1’s RFU. We measured the RFU from a sample of one hundred mutants from the epPCR library and found one mutant with significantly higher fluorescence than Mutant P1 (Figure 5A and B). Sequence analysis revealed a pseudorecombination/hybrid product at the C terminal end from two parental template, since *Taq* polymerase has a tendency to jump between templates during PCR amplification (Figure S4). Together, our directed evolution experiments using epPCR to enhance fitness of an artificially selected phenotype for pink fluorescent protein, resulted in a 20% increase in fluorescence from the first round, and a 35% increase in the second round (Figure 5C). The efficiency, as measured by the number of clones with increased fluorescence, decreased from 4 (out of 100) in the first round to 1 (out of 100) in the second round.

**Figure 5:**
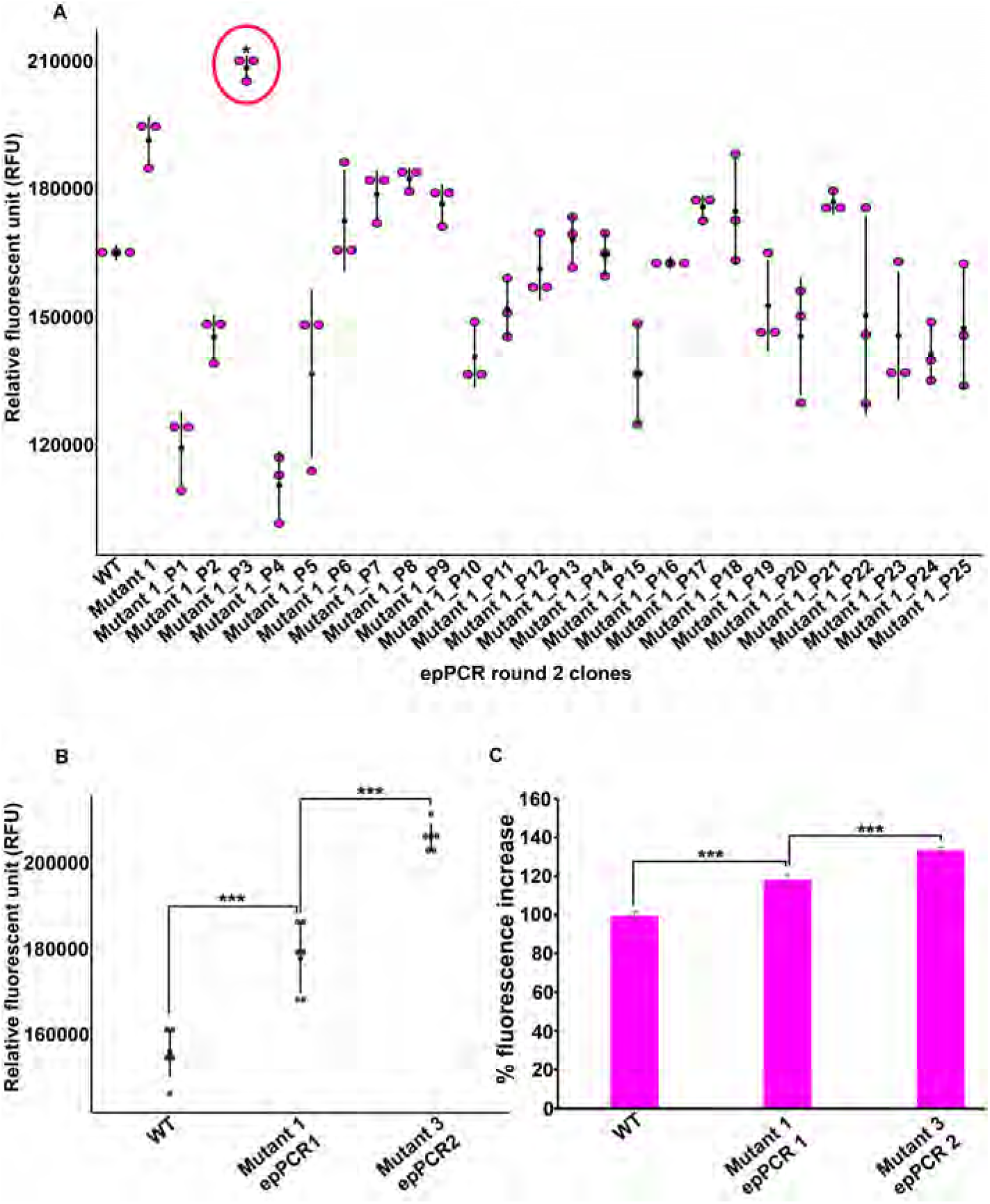
Iterative rounds of epPCR improved the fitness. **A.** RFU of 25 neutral mutants generated from second round of epPCR using plasmid from Mutant 1 (epPCR round1) as template. A mutant that displayed significantly increased fluorescence is shown in red circle (P represent pink colonies) (*, p*<*0.05). **B.** Dot plot showing increase in fluorescence between Mutant 1 (epPCR round 1) and Mutant 3 (epPCR round 2) in comparison to WT. **C.** Graph showing percentage increase in fluorescence achieved from two rounds of epPCR (***, p*<*0.0005).

### 3.5 Iterative rounds of EASME improved discovery

In our demonstration of EASME, presented in this paper, we started using a simple EA to search the protein space. We randomly mutated amino acid sequences by deleting, inserting and substituting amino acids, selecting insertions and substitutions uniformly among the 20 standard amino acids. This strategy is similar to the base algorithm used in DE. Fitness was evaluated by comparing the mutated sequence to approximately 200 protein sequences similar to Magenta (acquired through a BLAST search), with homology to those labeled “fluorescent” increasing fitness, while homology to those labeled “non-fluorescent” decreasing fitness. Experimental evaluation of the 15 most promising candidates did not show any increased fluorescence (Figure 6A).

Posthoc analysis of the candidate mutated sequences revealed that they contained a large number of mutations — an average of 56.33 mutations across all fifteen variants. This motivated using, in a second round of sequence exploration, a more conserved mutation function, mutating only set numbers of amino acids and then never modifying an already-mutated variant again. We tested mutation rates of one, three, or five amino acids, by sampling without replacement the mutation positions from the region of interest, and substituting the amino acid with a randomly-chosen one. We also modified the fitness evaluation algorithm by adding two machine learning models to the previous homology-based model: the first model estimating the propensity for protein aggregation and the second model estimated protein structural stability. These models were trained on thousands of input proteins with pre-defined fitness values. Because the evaluation of protein stability is computationally expensive, we used the homologybased fitness evaluation as a filter to identify promising candidates for further evaluation by these machine learning models. Using this algorithm, we identified 15 mutant with high homology and high predicted values from the machine learning models. Thus the models were executed serially to triage candidates with least computationally expensive implementations preceding others. In our experimental validation, laboratory RFU measurements revealed that 9 (out of 15, 60%) of the predicted variants were nonfunctional, being non-fluorescent, while 6 (out of 15, 40%) retained mildly pink phenotype (Figure 6B). None showed increased fluorescence.

In the third round, we further constrained the search algorithm by allowing only one mutation. We used the same homology-based filter, favoring sequences aligned more closely to proteins exhibiting fluorescence at the target excitation and emission frequencies. Fitness was evaluated by the same functions machine learning models used in round two. Fluorescence for the fifteen most promising candidates were evaluated experimentally. Of them, 11 (out of 15, 73%) mutants retained intermediate pink fluorescence, Figure 6C). While the observed increase in the number of fluorescent mutants is encouraging, the p-value from a 2-sample test for equality of proportions with continuity correction, using a one-sided alternative, is 0.07 (from prop.test() in R). Thus, we observed a jump of generated clones with intermediate fluorescence from 40% in the second round to 73.3% in the third. Similarly, the percentage of non-functional variants dropped from 60% in the second round to 26.7%. These results support the hypothesis that we could tune the algorithm, but not yet produce increased RFUs. We conclude that the EASME software package will improve with further research and optimized implementations (discussed below).

## 4 Discussion

A limitation of protein design is the enormous sequence space of any given protein. The protein search space is a vast mathematical space encoded by 20*^L^* (*L* represents sequence length) possible amino acid combinations for proteins of fixed length *L*, and even larger if we allow for indels. While DNA sequence databases continue to grow exponentially in size, for example GenBank currently contains over 3.44B DNA base pairs [59], their size are dwarfed by the size of all possible DNA sequences, leaving out a large fraction of potential DNA sequences from the available records as well as being biased towards extant sequences. Therefore, there will always remain a need for DNA evolutionary models to effectively search DNA space and predict the effect of mutations. In this study, we used modern directed evolution strategies to train a novel protein engineering paradigm, EASME, which combines EAs with fitness models [38].

We investigated this strategy by empirically evaluating candidate proteins our algorithm suggested as having improved functionality. To benchmark natural evolution against EASME, we first investigated the effectiveness of random mutation [60] (i.e., substitution of one amino acid at randomly chosen site) to enhance fitness. We experimentally introduced random point mutation at ten random sites, and the variants were laboratory tested for fluorescence. Of the ten variants tested, 7 (out of 10, 70%) mutants harbored deleterious or catastrophic mutations yielding non-functional or mildly functional pMagenta protein. While 3 (out of 10, 30%) variants displayed a neutral phenotype (Figure 1D and E). We concluded that simple random search was inefficient for producing variants with enhanced fitness, but this information is important to train the model and inform on the fitness landscape, as this approach dissects rates and placement of deleterious and catastrophic mutations [61, 62]. Our next approach used UV mutagenesis to generate variants. While UV mutagenesis accelerates the natural mutation rate, the mutations occur throughout the entire genome, making this approach inefficient for engineering specific proteins. To wit, the fluorescence of all the tested UV mutants was less than that of the wild-type.

Next, we employed epPCR to enhance fluorescence by creating random mutant library scanning the sequence space for beneficial mutations. In distribution of fitness effect (DFE) studies, researchers map the effects of all possible amino acid changes in a given protein to estimate rates of beneficial, deleterious, and neutral mutations [63, 64, 65]. These measurements inform on fitness landscapes that define protein engineering constraints for a given protein [66, 67]. DFE analyses reveal *≈*80% of mutations are deleterious [68]. Both random site-directed mutagenesis and epPCR results concurred with these observations (Figure 3B); we found that only about 0.8% of the total variants were beneficial and thereby improve an artificially-selected phenotype.

As our primary aim was to map beneficial mutations responsible for increased fitness, we initially generated a random mutant library with low mutation rate reflecting the idea that fewer mutations would result in higher amounts of enhanced function proteins. Sequence analysis revealed protein alleles that displayed enhanced fitness harbored two mutations on average. In terms of dynamic mutation rate optimizations, previous researchers who generated 15-30 mutations per gene [45, 69, 46, 44] observed that as the number of mutations increase in the gene, the fraction of proteins retaining the function declined exponentially. In contrast, high mutation rates result in lower numbers of functional sequences but can unlock novel regions of the search space and produce unique sequences. The success of protein engineering relies on generating a small library with high enrichment of functional proteins alleles [70]. Although we were able to achieve improved fitness using the epPCR approach, this method follows the hill climbing strategy of directed evolution, and as a consequence, this approach is often prone to local maxima preventing efficient navigation of the search space and the generation of novel sequences [71]. Multiple iterative rounds of epPCR are necessary to scan the sequence space and navigate fitness basins. EASME is a novel prototype AI. Herein we benchmarked the current EASME performance against modern epPCR methodology and were able to improve the algorithm’s performance moderately, but we were unable to generate an enhanced fluorophore *in this study*. By training the algorithm using lab data we directly obtained, we were able to improve the performance of our model (Figure 6). In the future, we’ll continue to improve the EASME algorithm with this approach.

A major limitation of the model was that it generated an exponential amount of bad sequences. We are currently optimizing and testing a large triage filter to reduce the rate of false-positive functional sequences (Figure 7). Another possible limitation of the model was inaccuracy of the fitness functions encoding theoretical calculations of brightness for the sequence variants. Estimating the brightness of a fluorescent protein relies on both excitation co-efficiency and quantum yield which depends on dynamic collective interactions taking place between the chromophore and the surrounding amino acid residues [72]. Capturing complex higher-order dependencies solely from changes in the sequence was challenging.

**Figure 6:**
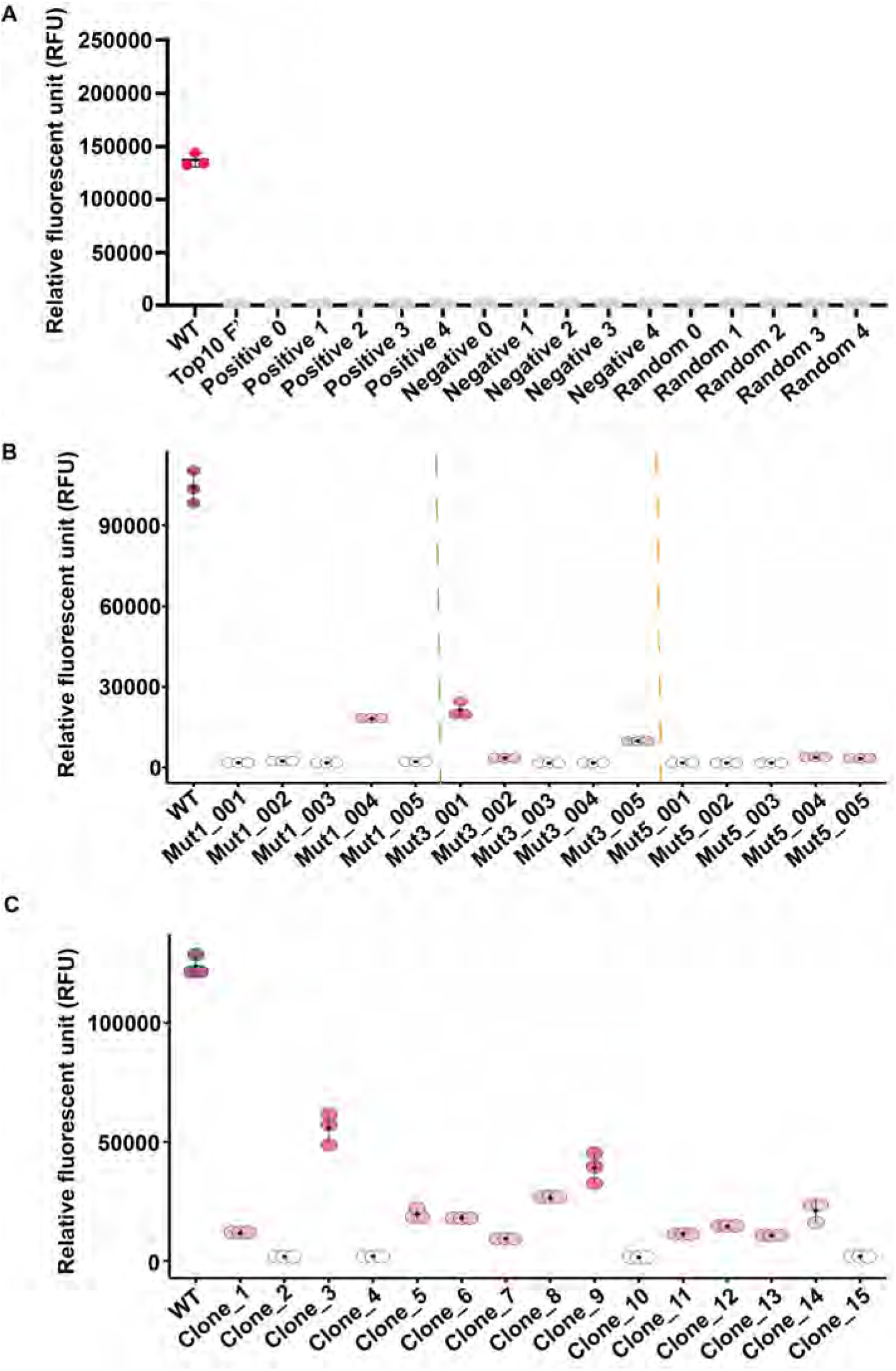
Iterative rounds of EASME tuned the algorithm and improved variant predictions. **A.** Dot plot showing RFU measurements from mutants generated in first round EASME. **B.** Second round of EASME. Mutants harboring single and three amino acid substitutions are separated by green and orange dashed lines respectively. The remaining five mutants posses five amino acid changes in Magenta. **C.** Third round of EASME. Magenta circle represent brighter pink fluorescence, pink circle represent mild fluorescence and white circle represent non-functional white variants.

**Figure 7:**
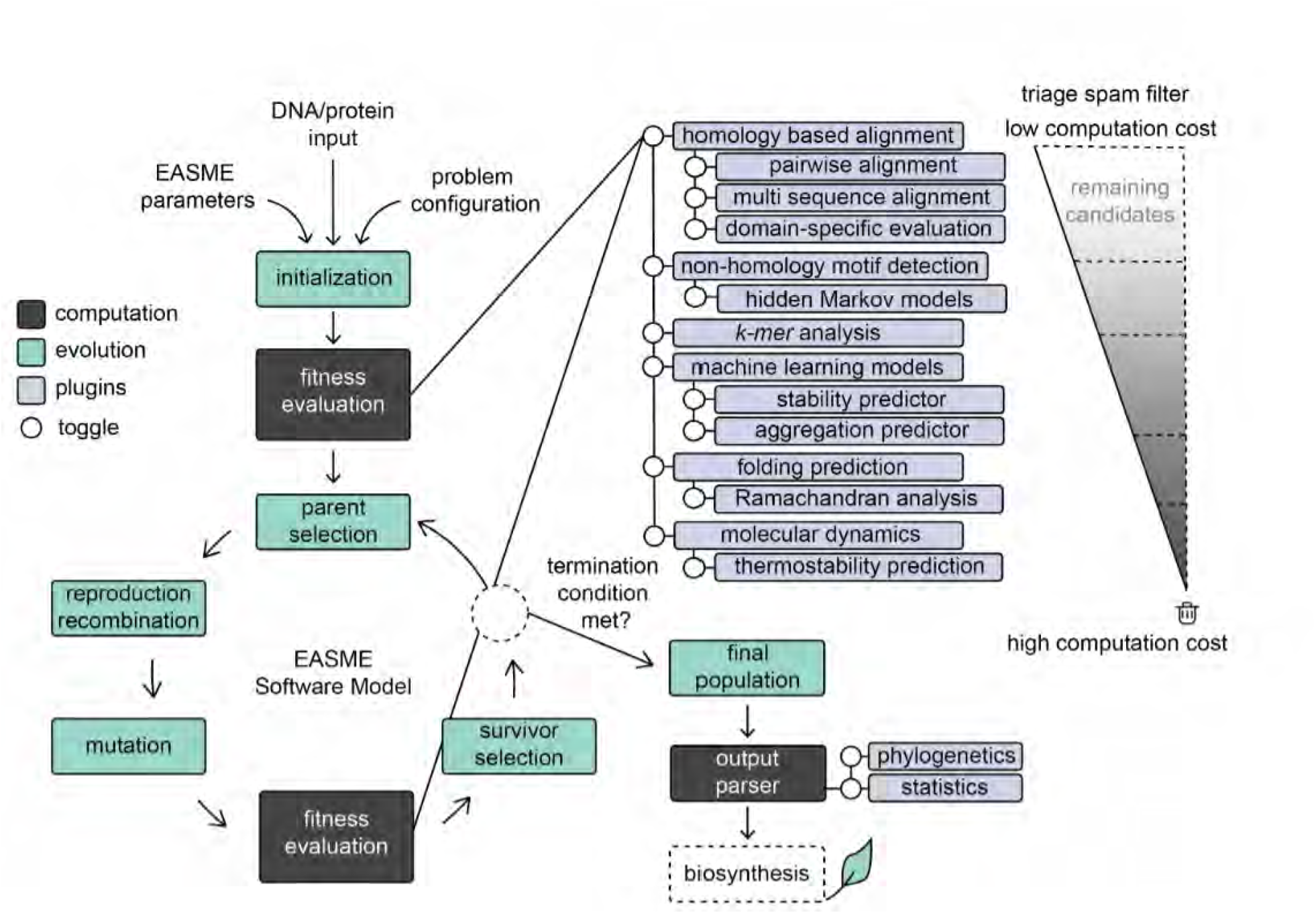
Schematic representation of a comprehensive model of the improved EASME toolkit indicating the different components of the model and fitness evaluation parameters.

The fundamental bottleneck of using current AI driven models for protein engineering is that these algorithms rely heavily on incomplete sequence libraries and only fine tune existing protein architecture rather than aiding in bold de novo re-design of proteins. The weakness of current AI models is evident when the algorithms only predict variants with slightly improved fitness such as a dim fluorescent protein, avGFP [73, 74] but none of the models create variants that outperform the brightness of currently available state-of-art fluorophores such as mNeoGreen [75] and StayGold [76]; these variants were discovered by generating multiple rounds of random mutagenized libraries using epPCR. Overall, studies indicate that more work is necessary to improve both molecular methods of protein engineering and AI design of proteins.

At a conceptual level, the long term goal is for EASME to improve both screening and scoring of variants. For now, epPCR beats EASME and probably most other AI based approaches in generation of true-positive enhanced fluorescent fluorophores (at least in our hands). But in proteins or enzymes where screening phenotypes is more complicated than visual evaluation of fluorescence, computational de novo AI engineering might be the only feasible path. Our long term goal is to develop an algorithm that (i) will effectively search the vast protein space, making smarter jumps to generate successful protein variants and (ii) more promptly and cost-effectively predict fitness functions of a given protein with higher levels of accuracy eliminating costly wet lab phenotype validations. The fact that we can improve EASME suggests that it shows promise as an alternative de novo protein design algorithm. Our longterm hypothesis is that EAs offer advantages over other forms of ML models, specifically in navigation of complex, jagged, and discontinuous search spaces [77]. To address this, we’re making meticulous changes to the current algorithm facilitating improved variant prediction (Figure 7).

In this study, we employed comparative benchmarking to explore hybrid strategies that combine AI with experimental approaches to increase the brightness of Magenta. Our results demonstrate that the EASME approach for protein design can be improved with mutational mapping (epPCR) studies that assist evaluating fitness functions through a feedback loop with wet-lab based results. One conclusion is that a suite of triage fitness functions needs to be developed, tested, and optimized to reduce the massive search space of protein exploration. One area of interest is concerned with *k*-mer algorithms that might efficiently and quickly crosscheck suggested proteins by cross referencing *k*-mers against a known biological lexicon. Thereafter, the logical path forward is to fine tune the EASME model with experimental datasets obtained from epPCR and toggling on/off a variety of proposed fitness functions that incorporate physicochemical structural information or physics-driven simulations that aid in defining chromophore specific regions, as previously suggested for GFP emission optimization [78]. We suggest these improvements will lead to a robust and complete EASME package and associated cyberinfrastructure, making this package accessible to scientists in biology, biotechnology, and related fields. Further, our study can serve as a reference point for future AI driven modeling studies in red fluorescent protein engineering, providing information to mark particular residues and regions.

## 5 Acknowledgment

We gratefully acknowledge funding for this work from the National Science Foundation via CAREER Award Number 2335888.

## A Supplementary Data

**Figure S1:**
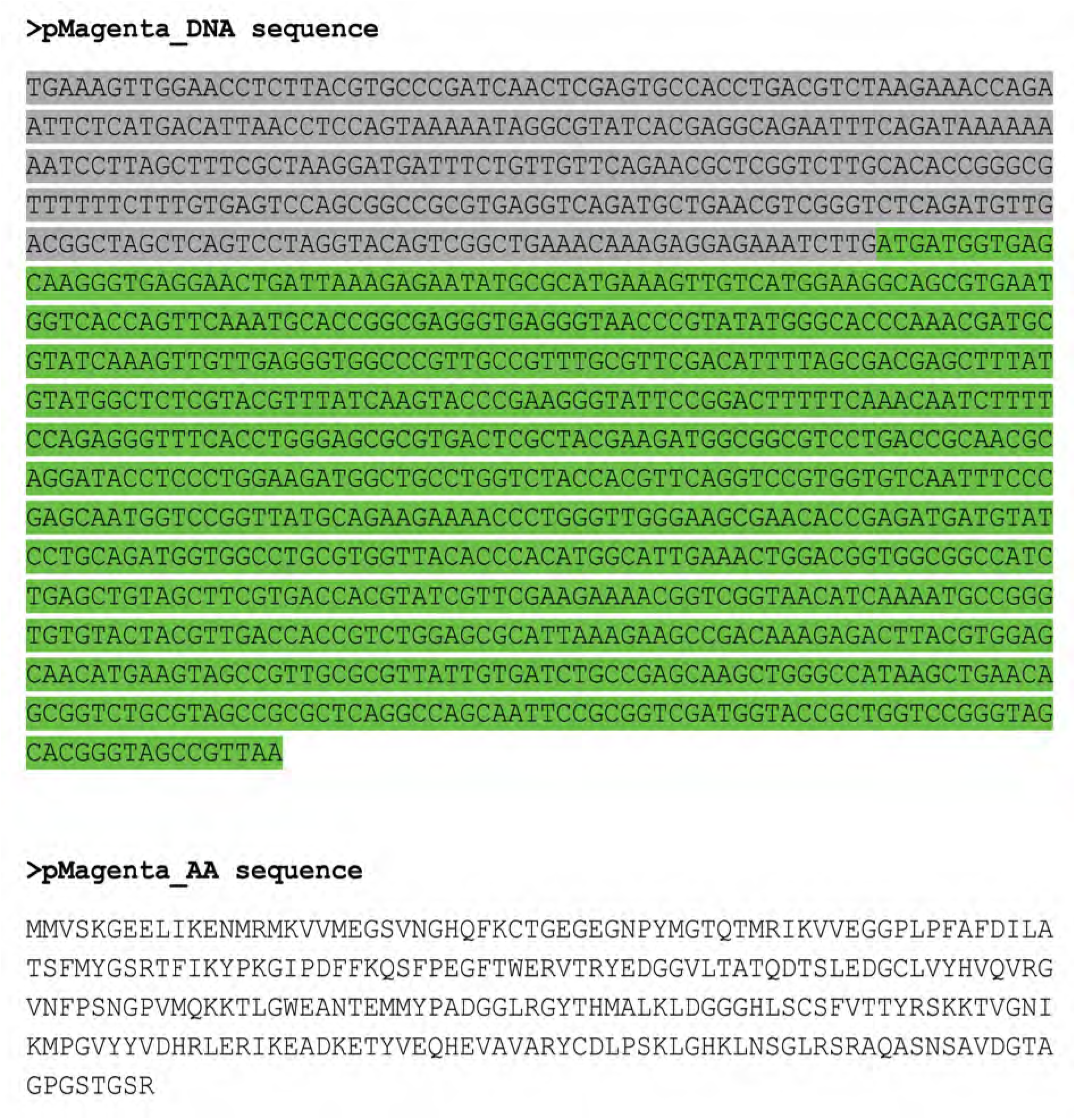
DNA and amino acid sequence of the pink fluorophore used in the present study. The region highlighted in grey represent the promoter and the coding sequence is highlighted in green.

**Figure S2:**
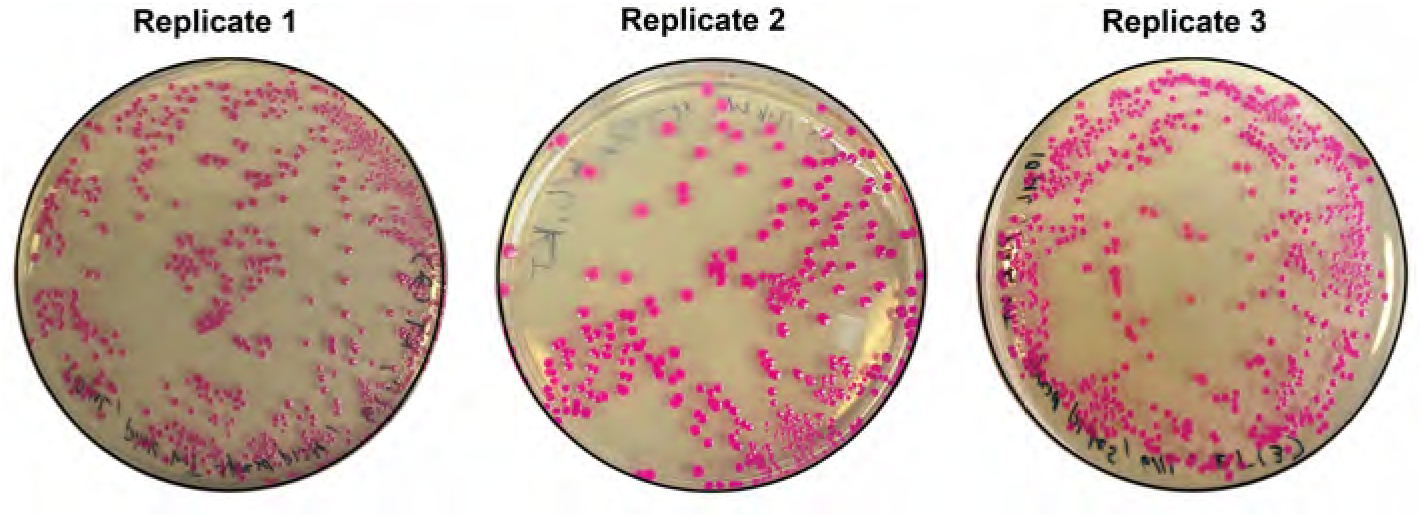
Accessing natural variation in *E. coli* expressing Magenta to discern neutral, deleterious and catastrophic mutations under optimal conditions. Representative plate image of *E.coli* harboring pMagenta plasmid cultured at optimal conditions for 48 hours. All colonies express fluorescence relative to wild-type suggesting no natural variation of *E.coli* cells. The experiment was performed in triplicate.

**Figure S3:**
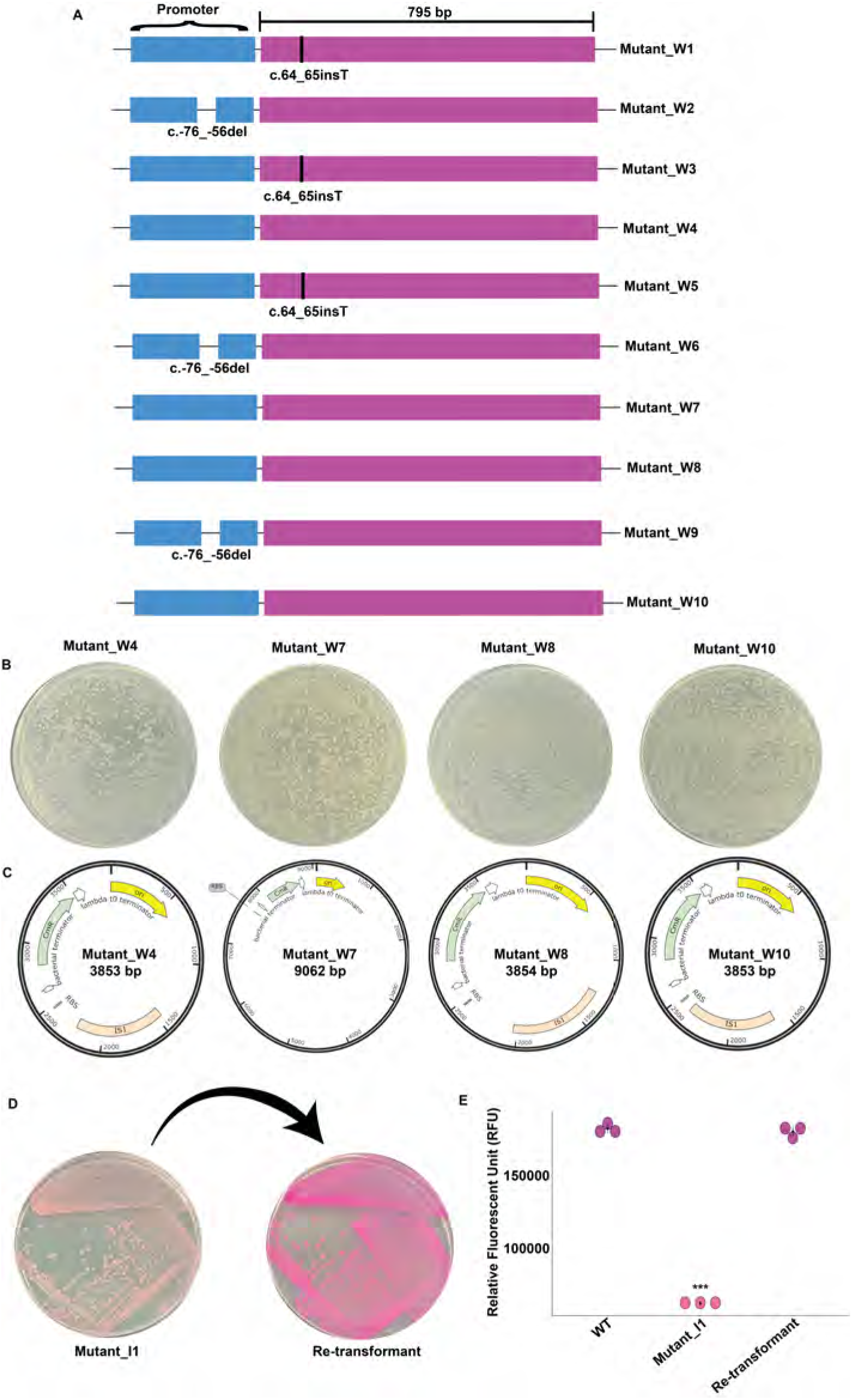
**A.** Mutation mapping indicating the position of indels identified in catastrophic mutants. **B.** Plates showing the presence of white colonies post re-transformation of plasmid extracted from catastrophic mutants W4, W7, W8 and W10. **C.** Plasmid map showing the integration of transposon elements in plasmids derived from mutant W4, W8 and W10 while plasmid multimerization in plasmid from mutant W7. **D.** Plate image showing the phenotype reversion from intermediate pink to wild type pink post re-transformation of plasmid from Mutant I1 to fresh *E. coli* Top10 F’ cells. **E.** RFU measurement of wild type, Mutant I1 and re-transformed clone (***, p*<*0.0005).

**Figure S4:**
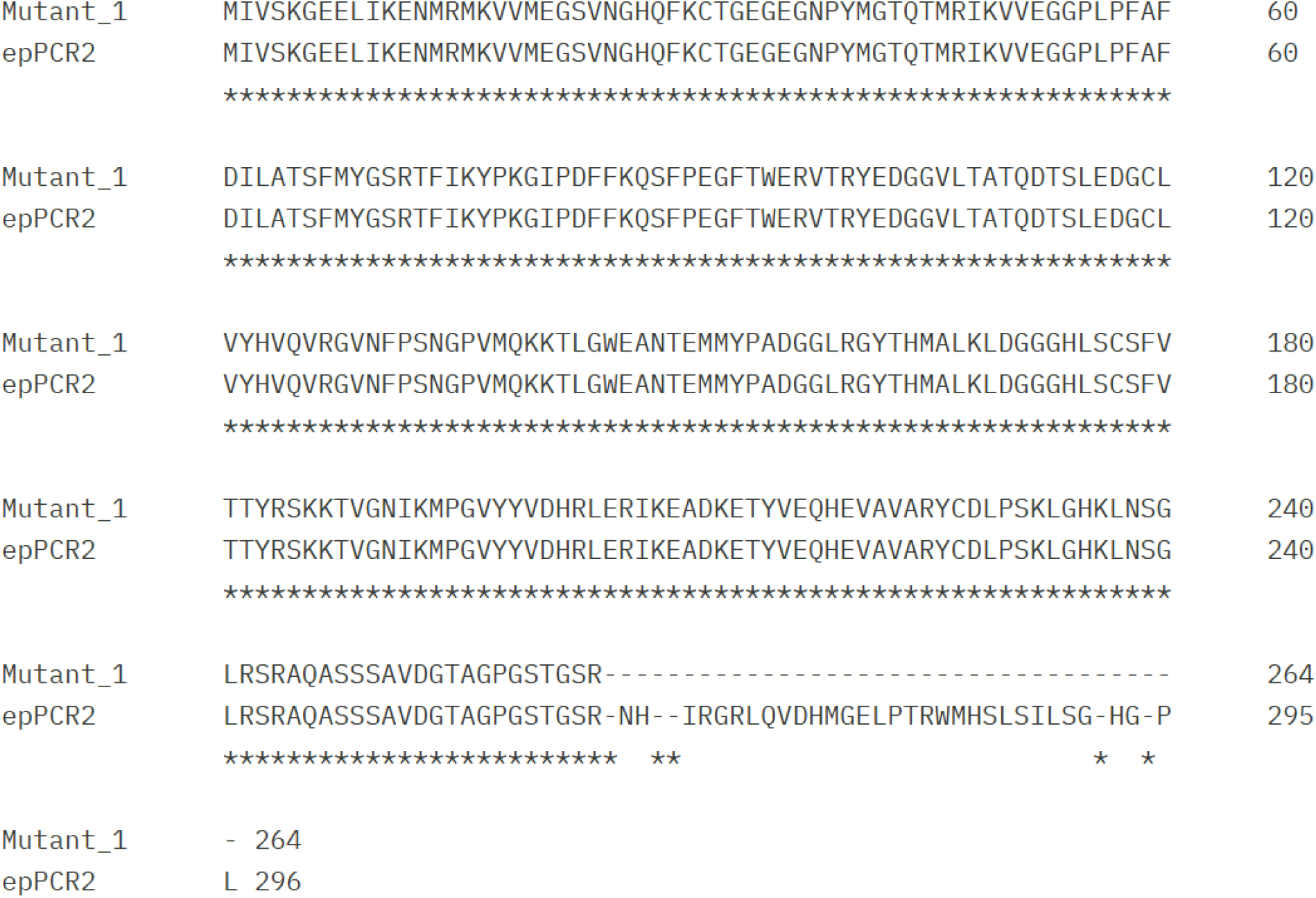
Alignment of Magenta coded by Mutant 1 (epPCR1) and Mutant 3(epPCR2) showing recombination at the C terminal end

**Table S1:** Bacterial strains and plasmids used in the study.

| Bacterial strain | Characteristics | Reference |
| --- | --- | --- |
| *Escherichia coli* |  |  |
| Top10 F' | F- mcrA (mrr-hsdRMS-mcrBC) 80lacZ M15 lacX74 recA1 ara 139 (ara-leu)7697 galU galK rpsL (StrR) endA1 nupG | Laboratory stock |
| **Plasmids** |  |  |
| pMagenta | Cm<sup>R</sup>, original plasmid harboring gene for pink fluorophore | Laboratory stock |
| pGEM-T | Am<sup>R</sup>, Conventional vector used for TA cloning used for mutant library preparation | Promega |

**Table S2:**
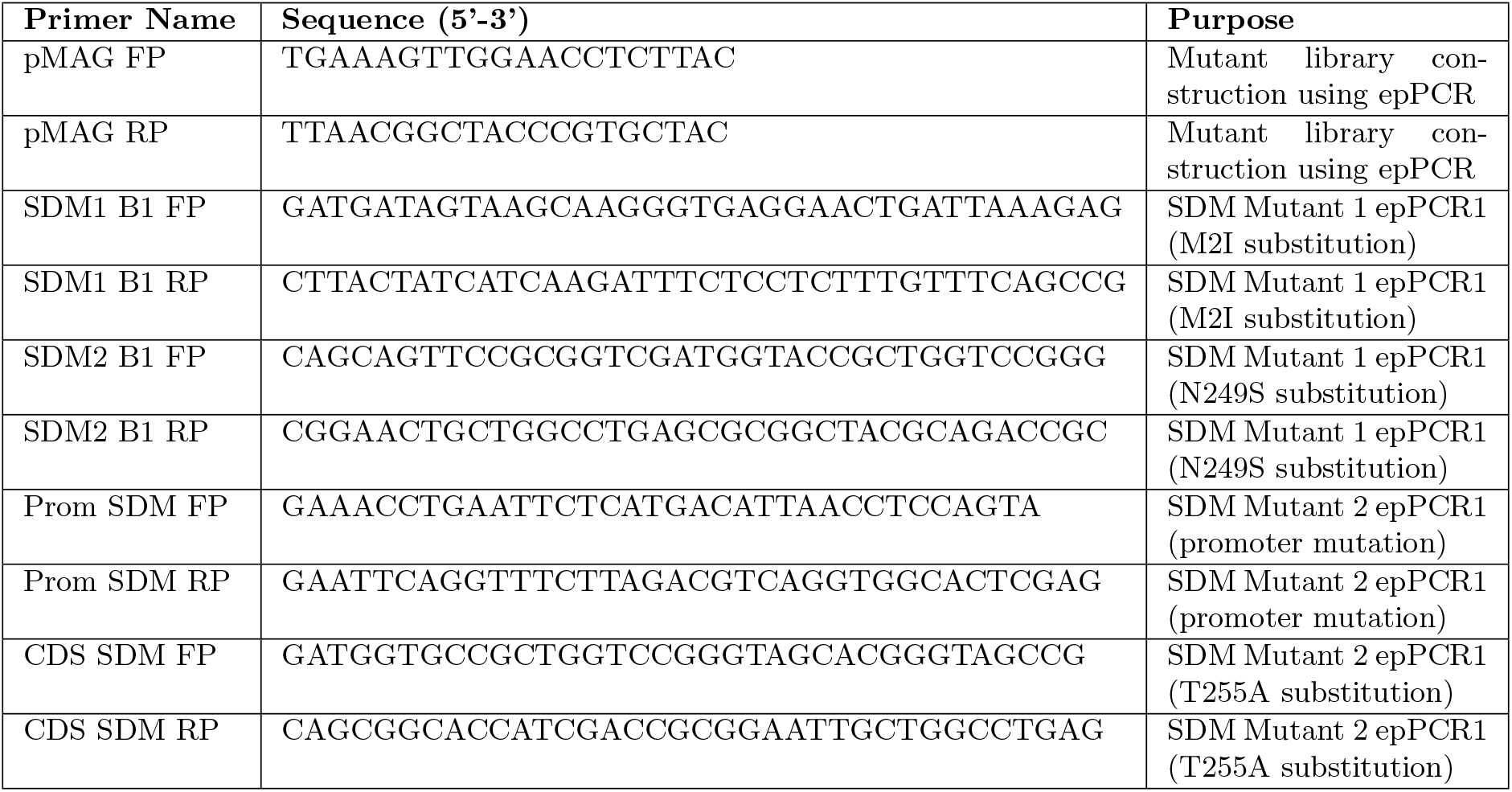
List of primers used in the study.

## Notes

### Competing Interest Statement

The authors have declared no competing interest.

